# Stimulus-induced narrow-band gamma oscillations in humans can be recorded using open-hardware low-cost EEG amplifier

**DOI:** 10.1101/2021.11.16.468841

**Authors:** Srividya Pattisapu, Supratim Ray

## Abstract

Stimulus-induced narrow-band gamma oscillations (30–70 Hz) in human electro - encephalograph (EEG) have been linked to attentional and memory mechanisms and are abnormal in mental health conditions such as autism, schizophrenia and Alzheimer’s Disease. This suggests that gamma oscillations could be valuable both as a research tool and an inexpensive, non-invasive biomarker for disease evaluation. However, since the absolute power in EEG decreases rapidly with increasing frequency following a “1*/f*” power law, and the gamma band includes line noise frequency, these oscillations are highly susceptible to instrument noise. Previous studies that recorded stimulus-induced gamma oscillations used expensive research-grade EEG amplifiers to address this issue. While low-cost EEG amplifiers have become popular in Brain Computer Interface applications that mainly rely on low-frequency oscillations (< 30 Hz) or steady-state-visually-evoked-potentials, whether they can also be used to measure stimulus-induced gamma oscillations is unknown. We recorded EEG signals using a low-cost, open-source amplifier (OpenBCI) and a traditional, research-grade amplifier (Brain Products GmbH) in male (*N* = 6) and female (*N* = 5) subjects (22–29 years) while they viewed full-screen static gratings that are known to induce gamma oscillations. OpenBCI recordings showed gamma response in almost all the subjects who showed a gamma response in Brain Products recordings, and the spectral and temporal profiles of these responses in alpha (8–13 Hz) and gamma bands were highly correlated between OpenBCI and Brain Products recordings. These results suggest that low-cost amplifiers can potentially be used in stimulus induced gamma response detection, making its research, and application in medicine more accessible.

## Introduction

Gamma rhythms are narrow-band oscillations in the 30-70 Hz range of the brain’s electrical activity [1]. They are associated with higher cognitive processes like attention [2, 3, 4], working memory [5] and feature binding [6], and are also found to be abnormal in mental health conditions like schizophrenia [7, 8], autism [9] and Alzheimer’s Disease (AD) [10]. Gamma oscillations can be induced in the occipital region of the brain when appropriate visual stimuli such as bars and gratings are presented to the subjects [11, 12], and such stimulus-induced gamma oscillations have been shown to decrease with healthy ageing [13] and with onset of AD [14]. Further, some studies have suggested a neuroprotective effect of entraining brain oscillations in the gamma range using sensory stimuli in rodent models of AD [15, 16, 17].

Power of EEG signals falls rapidly with frequency, following a 1*/f* power-law distribution [18]. Therefore, higher frequencies have much lower absolute power and are more susceptible to instrument noise. Mains (line) noise (50 or 60 Hz depending on the local power-line frequency) also lies within the gamma range. These factors make detection of gamma band oscillations difficult. These issues can be partially addressed by using research-grade amplifiers which have low input referred noise (noise generated by the internal circuitry of the amplifier in the absence of signals), high Common Mode Rejection Ratio (which amplifies the differential voltage while attenuating the common voltage between the positive and negative inputs), high input impedance to minimise the effect of high electrode impedance, and proper shielding of the electrical components of the amplifiers to reduce electromagnetic interference [19]. However, such amplifiers are generally bulky, expensive and often require proprietary software for usage. Several low-cost EEG acquisition systems have been developed in recent years (e.g. Emotiv (Emotiv Inc., San Francisco, California, U.S.A), NeuroSky TGAM, OpenBCI (OpenBCI, Inc., Brooklyn, New York, USA, www.openbci.com)), and their signal quality and performance has been studied in various contexts such as P-300 spelling task [20, 21], Motor imagery based BCI paradigm [22], drowsiness detection [23], motor tasks [24] and Steady State Visually Evoked Potentials (SSVEP) [25]. However, these studies involve assessment of signals present at frequencies lower than 40Hz. To the best of our knowledge, no study has tested whether stimulus-induced narrowband gamma rhythms can be detected using low-cost EEG amplifier systems.

We assessed the performance of OpenBCI, a popular affordable amplifier which provides a good cost-effectiveness [26], in the detection of gamma rhythms when static full-field gratings known to elicit gamma rhythms [12] were presented to healthy human subjects. OpenBCI recordings were compared to the recordings obtained using Brain Products GmbH, a popular research-grade amplifier, under identical experimental conditions on the same subjects. The same electrode cap, stimulus presentation software, and downstream analyses were used in both cases so that any difference was attributable only to the amplifiers. Full-screen gratings induce two distinct bands in the gamma range [12], termed slow gamma (20–34 Hz) and fast gamma (35–66 Hz) bands, which have characteristic spectral and temporal profiles [27]. Therefore, in addition to comparing the amplitude of change in band powers between stimulus and baseline, we also compared the spectral pattern and temporal evolution of the gamma bands between the two recording systems. In addition, we also compared the alpha band (8–13 Hz) power and temporal profiles.

## Methods

### Subjects

Eleven human subjects (6 males, 5 females, aged between 22–29 years) were recruited from the student community of the Indian Institute of Science, Bengaluru for the study on a voluntary basis against monetary compensation. Informed consent was obtained from all the subjects prior to performing the experiment. All procedures were approved by the Institute Human Ethics Committee of the Indian Institute of Science.

### Data Acquisition

For every subject, EEG signals were recorded using two amplifiers: BrainAmp DC EEG acquisition system (Brain Products GmbH, Gilching, Germany) and OpenBCI Cyton Biosensing Board. For proper comparison, both were connected to the same electrode cap, OpenBCI EEG Electrode Cap, a 21-channel setup with sintered Ag/AgCl electrodes (https://docs.openbci.com/AddOns/Headwear/ElectrodeCap/). We used 8 of these passive, gel-based electrodes at the following locations using the internationally recognised 10–20 system [28]: O1, O2, P7, P3, Pz, P4, P8, CPz. During acquisition, the EEG signals were referenced to Cz (unipolar reference scheme [29]). If the impedance of any electrode exceeded 25 kΩ, it was rejected offline during analysis.

In the Brain Products setup, raw signals were recorded at 5 kHz native sampling rate in AC coupled mode, filtered online between 0.016 Hz (passive R-C hardware filter) and 250 Hz (fifth-order low pass, Butterworth hardware filter) and digitised at 16-bit resolution (0.1 *µ*V/bit). Next, following an automatically applied digital low-pass Butterworth filter of 112.5 Hz cut-off to prevent aliasing, the data was downsampled to 250 Hz. This signal processing pipeline was implemented using BrainVision Recorder (Version 1.20.0701, Brain Products GmbH, Gilching, Germany). OpenBCI offers only an 8-channel recording with the requisite 250 Hz sampling rate, using a Bluetooth transmitter. While 16 channels can be used with an add-on board (OpenBCI Daisy board), it reduces the sampling rate to 125 Hz, which is too low for gamma range. A Wi-Fi shield was available which offered a higher sampling rate without losing on the channel availability, but it was still in beta phase at the time of our study. A higher sampling rate with 16 channels was also possible if data were recorded directly to the SD card on the equipment, but we opted to use streaming via Bluetooth for monitoring the signals in real time. For the OpenBCI setup, raw signals were recorded using OpenBCI GUI (version 5.0.2). Internally, OpenBCI first samples the signal at 1024 kHz in DC coupled mode followed by an R-C low-pass hardware filter of 72kHz. The signal is then digitised at 24-bit resolution (0.002235 *µ*V/bit) followed by noise-shaping and a digital, third-order, low-pass sinc filter as the anti-aliasing filter (https://www.ti.com/lit/ds/symlink/ads1299.pdf) before downsampling to our chosen sampling rate of 250Hz. It was observed during experimental setup that the OpenBCI system was sensitive to ambient mains noise, especially when the digital I/O pins were used to collect event marker data, and care had to be taken to prevent small perturbations from creating noise artefacts. To reduce line noise during acquisition, the OpenBCI setup was placed inside a copper mesh, grounded to the UPS ground socket, to serve as a Faraday cage. Eye tracking (monocular, left eye) was done for ten out of eleven subjects using Eye-Link 1000 (SR Research, Ontario, Canada) sampled at 1 kHz.

### Experimental Setting and behavioural task

All subjects sat in a dark room facing a gamma-corrected LCD monitor (BenQ XL2411; dimensions: 20.92 × 11.77 inches; resolution: 1289 × 720 pixels; refresh rate: 100 Hz) with their head supported by a chin rest at a distance of 57 cm from the screen. The experiment was performed in two sessions for each subject, one with OpenBCI and one with Brain Products (sequence chosen randomly) separated by a break for few minutes. Each session consisted of one minute of eye-open recording and one minute of eye-closed recording for measurement, followed by a visual fixation task. The entire experiment lasted for an average duration of 2.1 hours (minimum: 1.25 hours, maximum: 2.75 hours).

In the visual fixation task, each trial comprised of a 1 second fixation duration and 1 second stimulus duration, with a 1 second inter-trial interval. The stimuli presented were static, full-contrast, sinusoidal-luminance, achromatic gratings with a combination of one of the three spatial frequencies (1, 2 or 4 cycles per degree (cpd), calibrated to viewing distance) and one of the four orientations (0^°^, 45^°^, 90^°^ or 135^°^) and were displayed pseudorandomly using NIMH MonkeyLogic software (version 2.0.236 [30]). These stimulus parameters were chosen as they were previously shown to induce robust gamma [12]. Each of the two sessions consisted of an average of 298 ± 10 trials (mean ± SD), for the 12 stimulus types combined. Trials in which the eye tracker recorded an eye blink or a shift in eye position beyond a 5^°^ fixation window during fixation period or stimulus period were rejected online by the stimulus presentation software. The event markers for each stimulus type were conveyed to the two EEG acquisition devices using National Instruments USB-6008 Multifunction I/O Device.

### Artefact Rejection

A fully automated artefact rejection pipeline was used (for more details, see [13, 14]). Briefly, trials with deviation from the mean signal in either time or frequency domains by more than 6 standard deviations were labelled as outliers and rejected. Subsequently, data from electrodes with too many outliers (*>* 40%) was discarded. This resulted in a rejection of 21.4 ± 16.6% (mean ± SD) of trials for the OpenBCI session and 11.8 ± 7.2% of the trials for the Brain Products session. Finally, any electrode whose slope of the baseline power spectrum in the 56–84 Hz range was less than zero was rejected. This led to the rejection of 3 electrodes in 2 subjects, 1 electrode in 1 subject and no rejection in the remaining 8 subjects in OpenBCI recordings. In the Brain Products recordings, 2 electrodes were rejected in 1 subject, 1 electrode in 1 subject and no rejection in the remaining 9 subjects. If either of the electrodes of a bipolar pair (see below for details) was marked for rejection, the whole pair was removed from analysis.

### EEG Data Analysis

All analyses were performed using bipolar referencing scheme. Every electrode was re-referenced offline to its neighbour, yielding 5 bipolar electrode pairs (P7-O1, P3-O1, CPz-Pz, P4-O2, P8-O2) from the 8 unipolar electrodes. All the data analysis was done using custom codes written in MATLAB (The Mathworks Inc., 2021, version 9.10.0 (R2021a)). Brain Products data extraction included the usage of the ‘bva-io’ plug-in of EEGLAB toolbox (v12.0.2.5*b* [31], RRID: SCR 007292). Voltage measurements in OpenBCI recordings were sign-flipped [21] and were corrected offline to match the Brain Products standard while plotting Event Related Potentials (ERPs). The mains noise component was selectively attenuated offline prior to spectral analysis as follows [32]. First, we divided the unsegmented raw data into 180-second segments that contain integer cycles of the mains noise frequency. For each segment, we identified the frequency with maximum power using Fast-Fourier Transform in the 40–60 Hz range (to account for minor variations in the line noise frequency). A pure sinusoidal wave of that frequency (generated using inverse-FFT with the power of all other frequencies set to zero) was subtracted from the raw data to obtain the mains-noise-filtered data. Because the line noise component was notched out at high-frequency resolution, it was invisible in PSDs computed over short time segments. Linear detrending was done to the raw EEG signals to correct for slow drifts. Power Spectral Densities (PSDs) and the time–frequency spectra were computed using the multitaper method [33] with a single taper using the Chronux toolbox (http://chronux.org/, RRID:SCR 005547 [34]). With timepoint 0 marking the onset of stimulus, baseline period was chosen between *−*750 ms and 0 ms, and stimulus period was chosen between +250 ms and +1000 ms, yielding a frequency resolution of 1.33 Hz for the PSDs. The periods were chosen to avoid stimulus-onset related transients. Time–frequency power spectra were calculated using a moving window of size 250 ms and a step size of 25 ms, giving a frequency resolution of 4 Hz.

Change in Power between stimulus and baseline periods for a frequency band was calculated using the following equation:

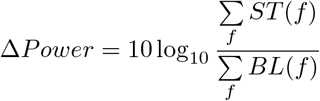

where *ST* is the stimulus power spectrum and *BL* is the baseline power spectrum, both averaged within relevant frequency bins (*f*), across all analysable trials and electrodes. Band powers were specifically computed in three frequency bands, namely slow gamma (20–34 Hz), fast gamma (35–66 Hz), and alpha (8–13 Hz).

### Slope of the PSD plot

For slope calculation, the PSDs were fit with the following power law function [18]:

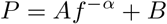

where *P* is the power and *f* is the frequency, while *A* (scaling function), *B* (noise floor), and *α* (slope) are free parameters. To avoid over-fitting, we set B as the power at max frequency (125 Hz). Subsequently, linear regression was done on the log of Power (after subtracting the noise-floor) to obtain an estimate of *A* and *α*. As in our previous paper [13], slopes were calculated specifically for the 16–34 Hz and 54–88 Hz ranges to avoid contamination by the alpha range bump and line noise artefact and its harmonics.

### Correlation Analysis

Similarity between OpenBCI and Brain Product recordings was quantified using the Spearman correlation of the data points between the two sessions. For band powers, the data points were the change in band power between the stimulus and baseline averaged across electrodes (yielding one value per frequency band for each subject). For assessment of similarity of temporal evolution, the data points were the time series of the mean band power change obtained using time frequency spectra. Correlation measures between OpenBCI and Brain Product recordings of the same subject are termed ‘self-pair’ correlations, and between OpenBCI and Brain Product recordings of two different subjects are termed ‘cross-pair’ correlations.

### Statistical Analysis

Appropriate non-parametric tests (Mann Whitney U test (Wilcoxin rank sum test) [35], permutation test [36]) were used to interpret the findings. 0.05 was used as the cut-off for significance of the p-values.

### Data and Code Availability

The data and codes used in this study are all made publicly available and can be found at https://github.com/FlyingFordAnglia/OpenBCIGammaProject.

## Results

### Instrument noise

Before EEG recordings, we characterized the internal instrument noise characteristics by placing the recording electrodes, along with the reference and ground in a common conducting salt bath (dashed lines in Fig 1). The power spectral density (PSD) for OpenBCI recordings (red dashed lines) showed larger line noise at 50 Hz compared to Brain Products (blue dashed lines), and also exhibited three additional peaks at 14 Hz, 36 (50 *−* 14) Hz and 64 (50 + 14) Hz. These peaks are due to modulation of the mains noise, that occurs due to non-linear distortion during amplification. No such artefactual peaks were observed for the Brain Products system. Even after shielding the OpenBCI setup for EEG recordings (which reduced the noise peak at 50 Hz, see Methods for details), these three peaks could be observed for some subjects (as shown below).

**Figure 1.**
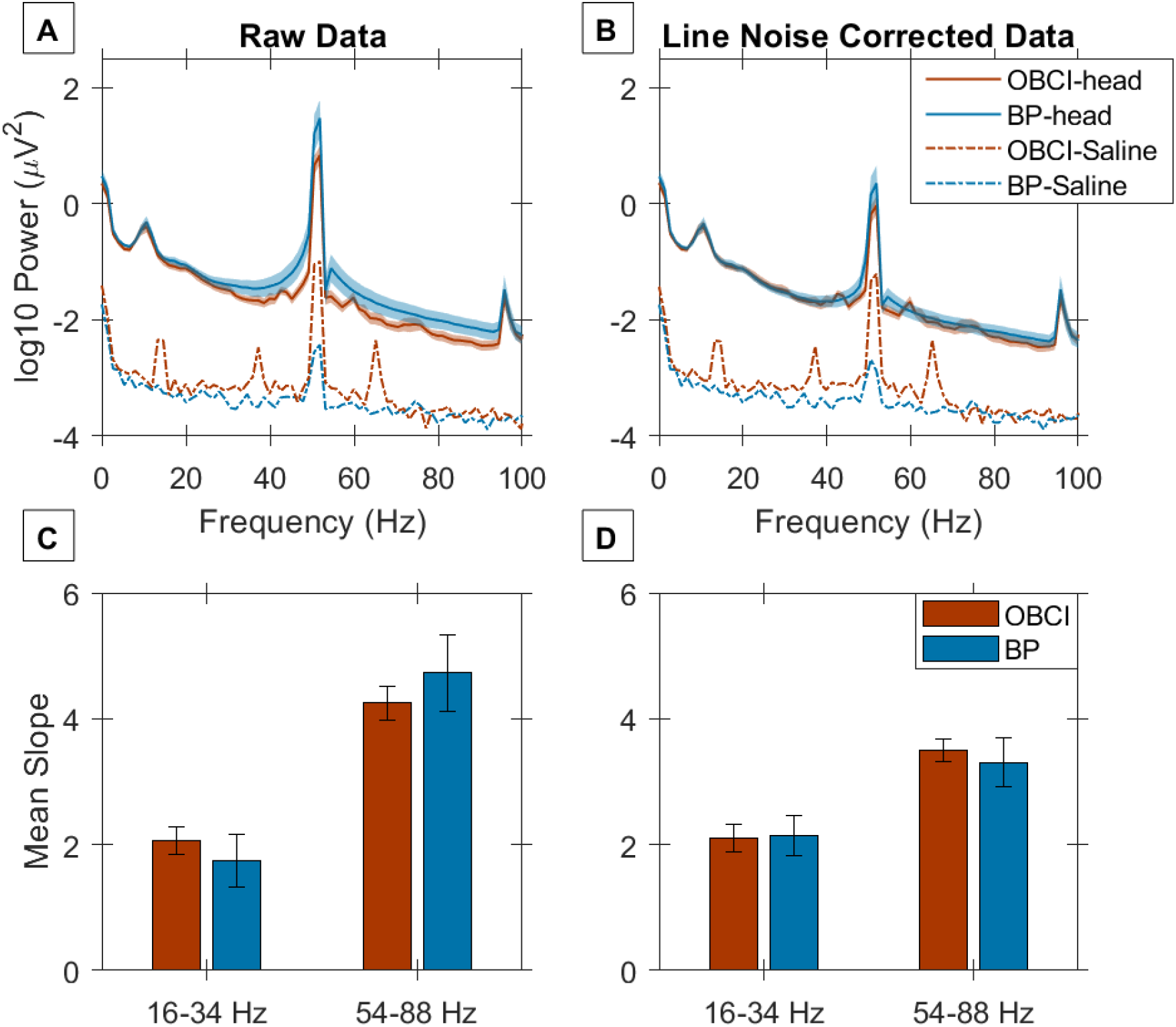
Slope comparison of OpenBCI and Brain Products at baseline. **A)** Baseline raw Power Spectral Density (PSD) for OpenBCI (*red trace*) and Brain Products (*blue trace*) averaged across 11 subjects (thickness of the trace indicates the standard error of PSD across subjects at each frequency). Dotted lines show PSD of shorted electrodes for instrument noise measurement. **C)** Slope of the PSD vs Frequency plot averaged across all subjects for two frequency ranges (16–34 Hz and 54–88 Hz) for OpenBCI (*red*) and Brain Products (*blue*). The error bars indicate the standard error. **B, D)** Same as A, C, but after selectively attenuating the mains/line noise component (see Methods for details).

### Baseline PSDs and Slopes are comparable

Next, we compared EEG recordings in the pre-stimulus baseline period. Fig 1A (solid lines) shows the mean PSD across all subjects after averaging across all trials and up to 5 bipolar electrodes for each subject (see Methods for details). The Brain Products system had more mains noise than OpenBCI, possibly due to the usage of the Faraday shield for OpenBCI (no shield was used for Brain Products). To reduce the line noise artefact, we estimated the mains noise component in long segments of data and subtracted the same [32] before re-computing the PSD (see Methods for details), which yielded similar PSDs using both amplifiers (Fig 1B). The slopes of the PSDs computed at two different frequency ranges (16–34 Hz and 54–88 Hz as per our previous report [13]) were not significantly different between OpenBCI and Brain Products recordings (Fig 1C and 1D: raw-data: 16–34 Hz: *p* = 0.69, 54–88 Hz: *p* = 0.33, noise-corrected data: 16–34 Hz: *p* = 0.90; 54–88 Hz: *p* = 0.74; two-tailed Mann Whitney U test). The PSDs of the two systems were highly correlated (spearman correlation coefficient of 0.91 for raw data (*p* < 10^−6^, calculated using permutation test) and 0.93 for noise corrected data (*p* < 10^−6^)).

Fig 2 shows the results of an example subject (subject S2) for the visual fixation task. Trial and electrode averaged evoked potentials are plotted for OpenBCI (Fig 2A, left column) and Brain Products (Fig 2A, right column). These traces revealed a transient in the first 200 ms after stimulus onset and after the stimulus offset (i.e. after +1000ms). Change in power in stimulus period compared to baseline power revealed a prominent suppression in the alpha range and in increase in slow gamma power in both OpenBCI and Brain Products (Fig 2A, second row). There was also a broadband increase in power beyond the slow-gamma range, which was more prominent in Brain Products compared to OpenBCI. Power increase in the alpha band after 1000ms of the trial was likely an eye blink or movement artefact during the inter-trial period. Fig 2C shows the PSD of the stimulus and baseline periods for the two amplifiers, demonstrating alpha (8–13 Hz) suppression and an increase in power in the slow-gamma (20–34 Hz) and the fast-gamma (35–66 Hz) bands, with the fast-gamma response being more appreciable in the Brain Products recording. Fig 2D shows the baseline subtracted PSDs for the two systems illustrating the same trend as above (since the log of PSD is subtracted, it is essentially a change in power from baseline, expressed in decibels). The change in power in alpha, slow and fast gamma bands as a function of time from their respective pre-stimulus baseline power is shown in Fig 2B. Overall, the increase in the band power is lower in OpenBCI than Brain Products for both the gamma bands, while it is similar for the alpha band (Fig 2A second row, 2B, 2C, 2D). Small noise peaks placed symmetrically around the mains noise band, indicating modulation distortions (also see Fig 1), can be observed in the PSDs of OpenBCI in this subject (Fig 2C, first column). However, since this noise is present in both pre-stimulus baseline period and stimulus-period, it gets cancelled out when we compute the change in power from baseline (Fig 2C, 2D).

**Figure 2.**
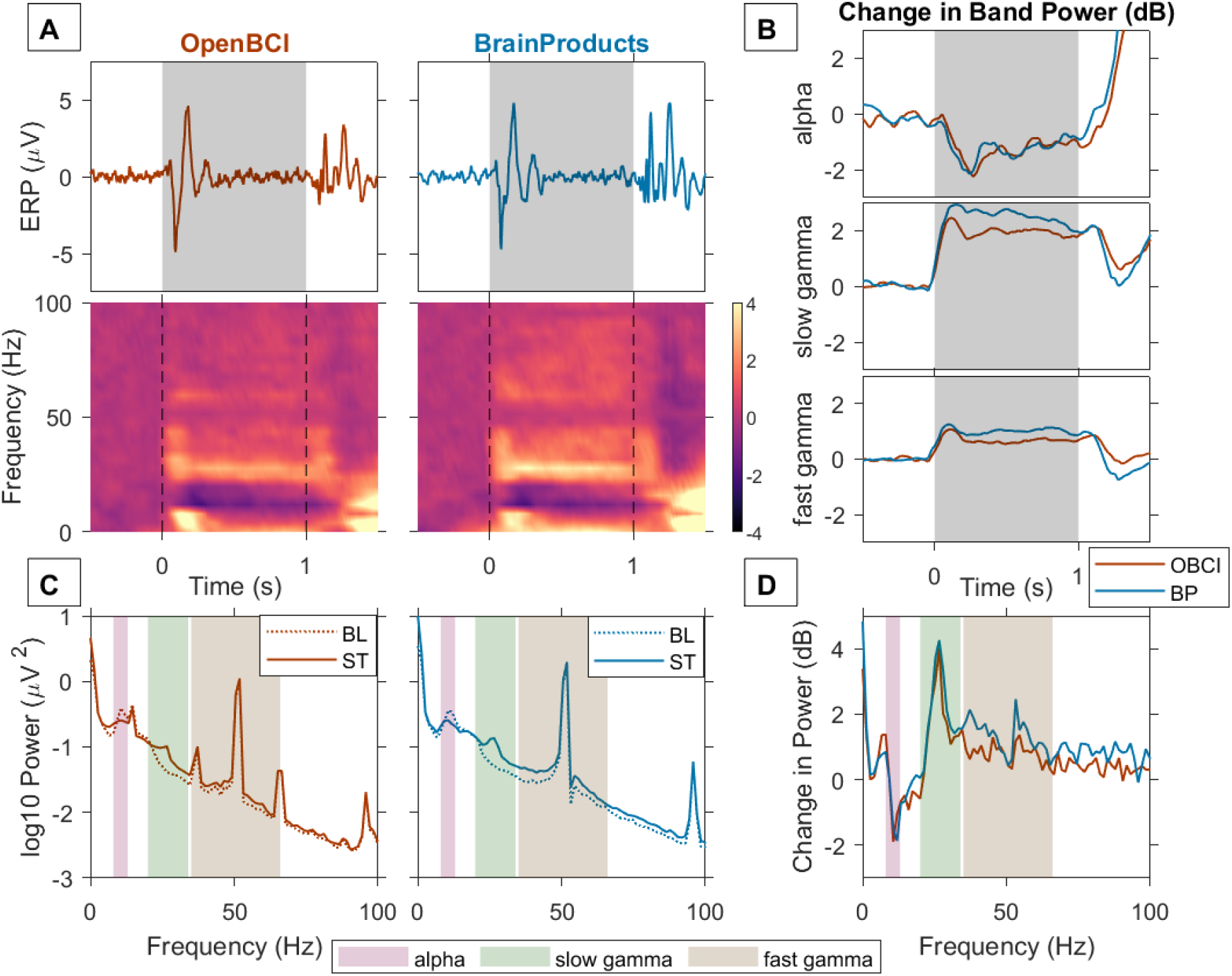
Stimulus induced gamma response in an example subject. **A)** Trial averaged EEG trace or ERP (*first row*) and time frequency spectrograms of change in power from baseline with time (*second row*). Vertical grey bars indicate stimulus duration (1 second). **B)** Change in Power (dB) from baseline as a function of time in alpha (8-13 Hz, *first row*), slow gamma (20–34 Hz, *second row*) and fast gamma (35–66 Hz, *third row*) bands recorded using OpenBCI (*red trace*) and Brain Products (*blue trace*). **C)** PSD for stimulus (*solid line*) and baseline (*dotted line*) averaged across 5 bipolar electrodes for OpenBCI (*left column*) and BrainProducts (*right column*). **D)** Change in power (dB) from baseline in stimulus period recorded using OpenBCI (*red trace*) and Brain Products (*blue trace*).

### Spectral and temporal patterns show similarity

Fig 3 shows the results of the visual fixation task for all the subjects sorted by decreasing gamma power. Visually similar results were obtained in the baseline-subtracted time frequency spectra of OpenBCI (first column) and Brain Products (second column). The change in power from baseline during stimulus at each frequency (Figure 3, third column), and the change in mean band power (in dB) of alpha, slow gamma and fast gamma bands with time (Fig 3, 4th, 5th and 6th columns respectively) also showed visually similar trends. However, the amplitude of change in band power can be seen to be lower in OpenBCI than in Brain Products in most subjects for the gamma bands, in particular for the fast gamma band.

**Figure 3.**
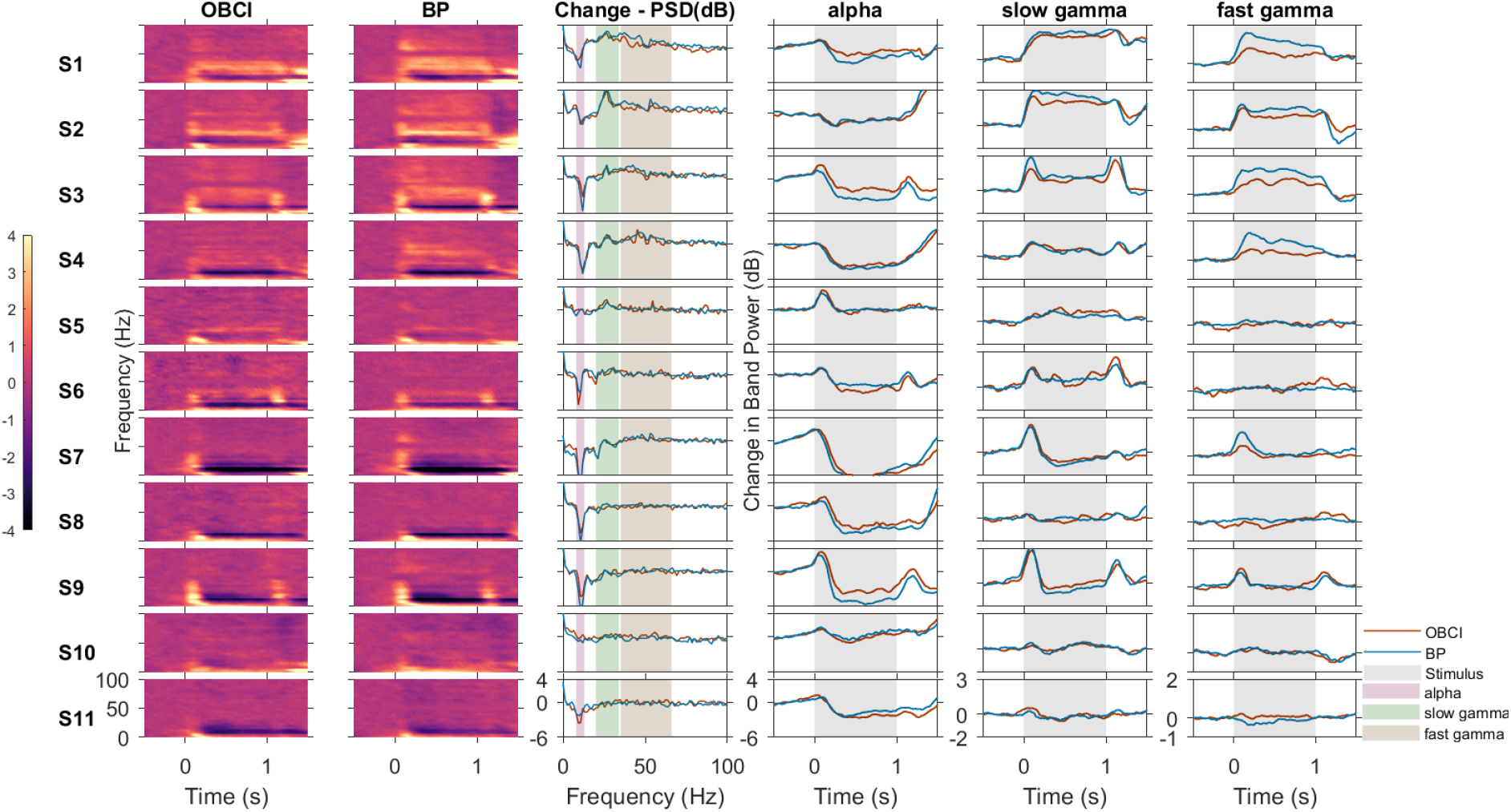
Comparison of OpenBCI and BrainProduct recordings for all subjects. Baseline subtracted time freqency spectrograms for OpenBCI (first column and *red trace* in other columns) and Brain Products (second column and *blue trace* in other columns), change in PSD (dB) from baseline in stimulus period vs frequency (third column), change in power (dB) with time for alpha (fourth column),slow gamma (fifth column) and fast gamma (sixth column) bands. Vertical bands in the last three columns indicate stimulus duration (*grey*). Each row represents one subject. The subjects are numbered in decreasing order of total gamma power.

We considered a subject as having a “gamma response” if they showed a significant increase in band power between stimulus and baseline when compared across all trials using one-tailed Mann-Whitney U test. False Discovery Rate (FDR) was controlled using the Benjamini and Hochberg procedure [37]. With this criterion, 6 subjects (S1-S6) were found to have slow gamma response using both Brain Products and OpenBCI. For fast gamma, 5 subjects (S1-S4,S7) showed a fast gamma response with Brain Products, out of which 4 subjects showed a fast gamma response (S1-S4) with OpenBCI. A significant alpha suppression was observed in 10 subjects (all subjects except S5) using Brain Products and 9 subjects (all subjects except S5 and S10) using OpenBCI. In subject S10, while it is not apparent visually in Fig 3, there was a very small fast gamma increase (*≈* 0.1 dB increase from baseline in stimulus period) in OpenBCI recordings that was registered as significant by the statistical test we used, even after multiple testing correction. Similarly, alpha suppression in this subject was significant with Brain Products (but not OpenBCI), even though the suppression was small (*≈* 0.76 dB reduction from baseline in stimulus period). Overall, these results indicate that the power changes obtained using Brain Products and OpenBCI were highly consistent. To quantify these results, we computed the correlation between the change in band power from baseline in each frequency band of each subject using OpenBCI with that of Brain Products (Fig 4). The Spearman correlation coefficient for alpha band was 0.92 (*p* < 10^−6^, computed using permutation test), for slow gamma was 0.94 (*p* < 10^−6^) and for fast gamma was 0.75 (*p* = 0.012). For frequency wise correlation values across all subjects, see Supplementary Fig S1.

**Figure 4.**
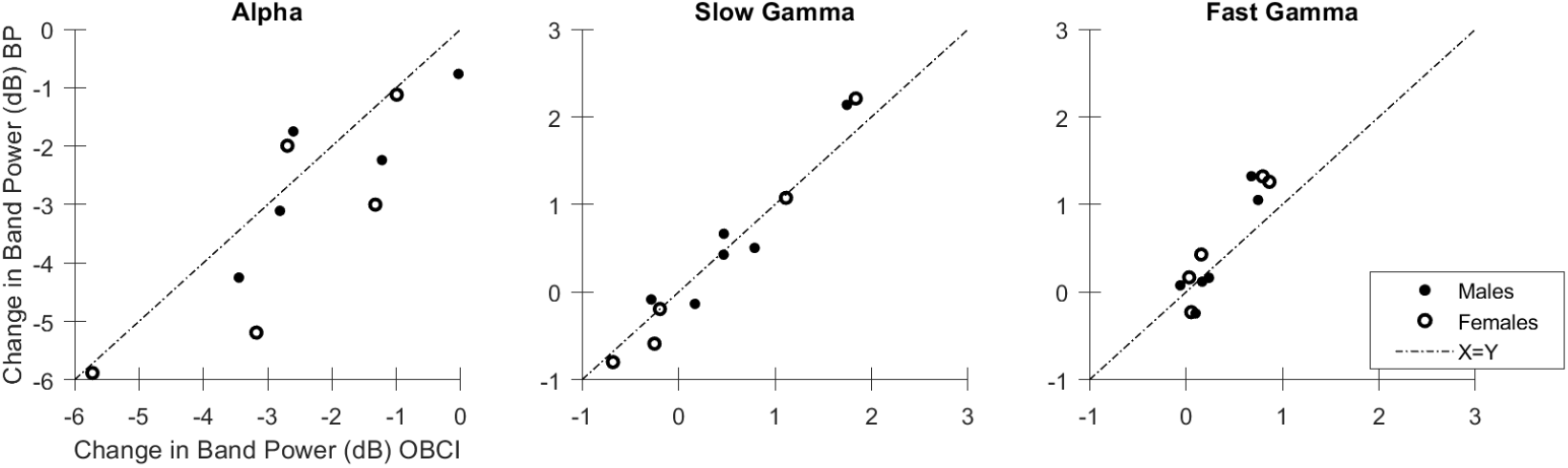
Comparison of Change in Band Powers of OpenBCI and Brain Product recordings. Change in band power from baseline in stimulus period of alpha, slow gamma and fast gamma oscillations recorded using OpenBCI (*x axis*) and Brain Products (*y axis*) for males (*filled circles*) and females (*open circles*). Dashed line indicates indicates identity line.

### Amplitude of change in band powers appears to be better captured by Brain Products than OpenBCI

Although the correlation between power obtained using the two amplifiers was high, the distribution of points around the identity line (Fig 4) indicates that the amplitude of band power change was generally greater in Brain Products than OpenBCI. For alpha band, most points lay below the identity line indicating that alpha suppression was better captured by Brain Products than OpenBCI. For fast gamma, the enhancement in gamma power was again better captured with Brain Products than OpenBCI, with the majority of data points above the identity line. For slow-gamma, the difference between the two devices appear less salient. However, note that the power in slow-gamma band is influenced by two opposing factors. First, there is an increase in power due to the slow-gamma rhythm, which is observed in about half of the subjects (S1-S6). However, there is also a reduction in power in lower frequencies, which sometimes extended to the slow-gamma range. This is better observed in the subjects (S7-S11) who did not have a strong slow-gamma rhythm, and their corresponding data points lay below zero dB in Figure 4 (middle panel). For those subjects, the points lay below the identity line, again reflecting better capture of low-frequency suppression by Brain Products than OpenBCI, and not poorer capture of the slow-gamma rhythm by Brain Products compared to OpenBCI.

### ERPs are comparable but with a small latency

Figure 5 shows the de-trended and de-noised (see Methods for details) ERPs of OpenBCI and Brain Products for all subjects. A slight jitter can be seen in the OpenBCI traces compared to the Brain Products traces. Computing the cross-correlation between the two traces led to a median correlation (± standard error; computed using bootstrapping) of 0.79 ± 0.05, with OpenBCI traces lagging by 8 ± 0.5 milliseconds (cross-correlation value and the lag for each subject are indicated in the figure).

**Figure 5.**
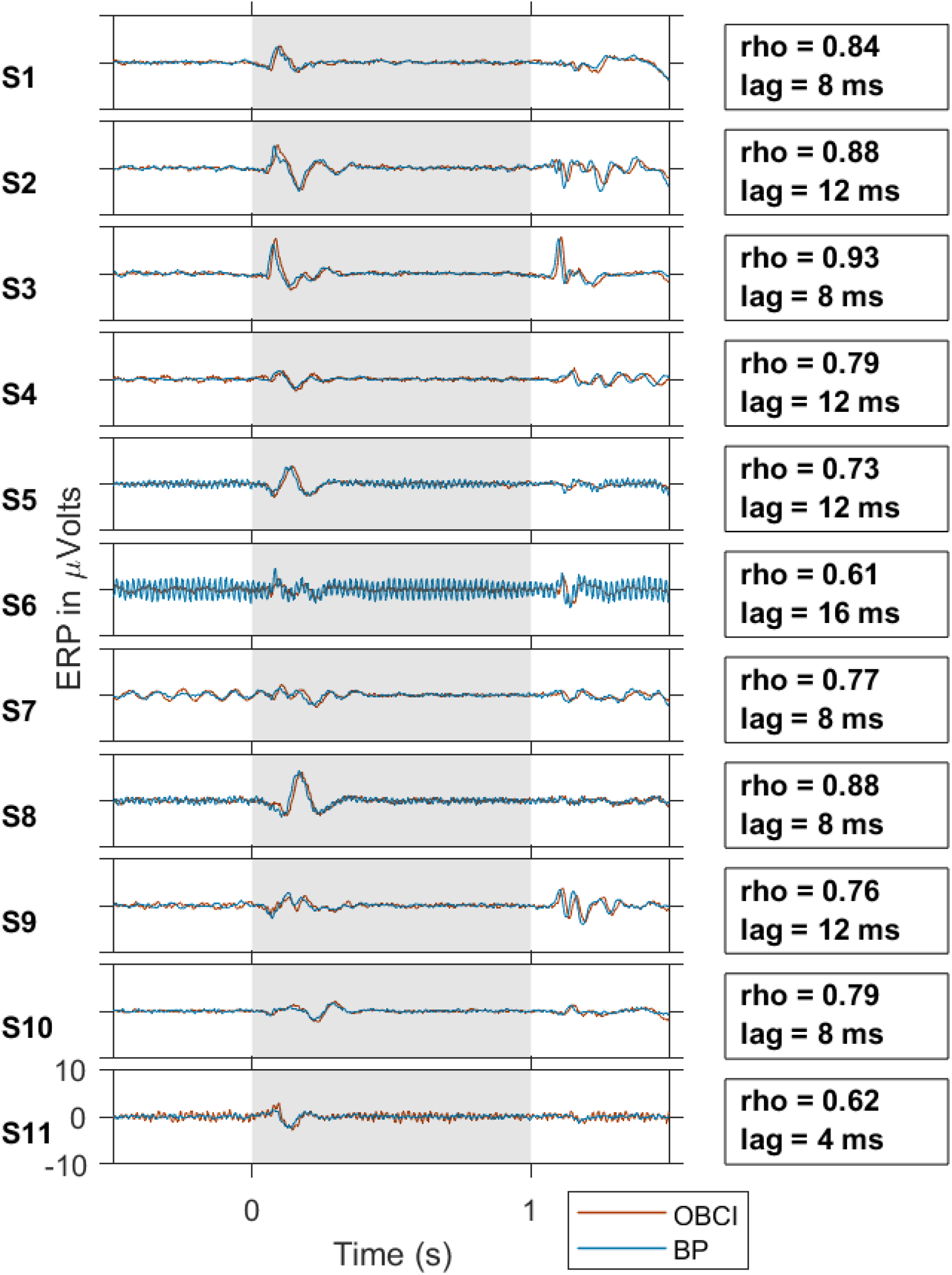
ERP Plots of all subjects. ERPs for OpenBCI (*red trace*) and Brain Products (*blue trace*) for all subjects. The subjects are numbered in decreasing order of gamma power, as in Figure 3. The vertical grey shaded region indicates the stimulus duration. The cross-correlation coefficient and lag between OpenBCI and Brain Products are indicated on the right for each subject.

### Similarity between the two recordings within and between subjects

Next, we compared the temporal profile and the characteristic spectral distribution (see Introduction) between the two setups. The similarity of the change in band powers with time, and baseline subtracted PSDs of the two EEG amplifiers was quantified using Spearman correlation between the time series for each subject (see Methods). However, a high correlation between OpenBCI and Brain Products traces may be confounded by the possibility that the spectral profile in response to visual stimulus is common to all subjects. A previous study suggests that spectral and temporal profiles are unique to subjects and different in different subjects [27]. To assert whether these correlations indeed represent subject-specific trends and not a general similarity of all traces, we further compared self-pair (between the same subject) and cross-pair (between different subjects) correlations. Self-pair correlations of baseline-subtracted PSD (0.51 ± 0.1, median ± standard error computed using bootstrapping) were significantly higher than its cross-pair correlations (0.33 ± 0.02, *p* = 1.07 × 10^−4^, one sided Mann Whitney U test, Fig 6A). Similar results were obtained for the temporal evolution traces in alpha, slow gamma and fast gamma bands:(alpha: self: 0.93 ± 0.04; cross: 0.52 ± 0.07, *p* = 7.4 × 10^−6^, Fig 6B; slow gamma: self: 0.82 ± 0.05; cross: 0.31 ± 0.06, *p* = 6.01 × 10^−6^, Fig 6C; fast gamma: self: 0.68 ± 0.15, cross: 0.22 ± 0.04, *p* = 2.7 × 10^−4^, Fig 6D). When self-pair correlations were plotted against cross-pair correlations, their values were concentrated on the x-axis side of the identity line (Fig 6).

**Figure 6.**
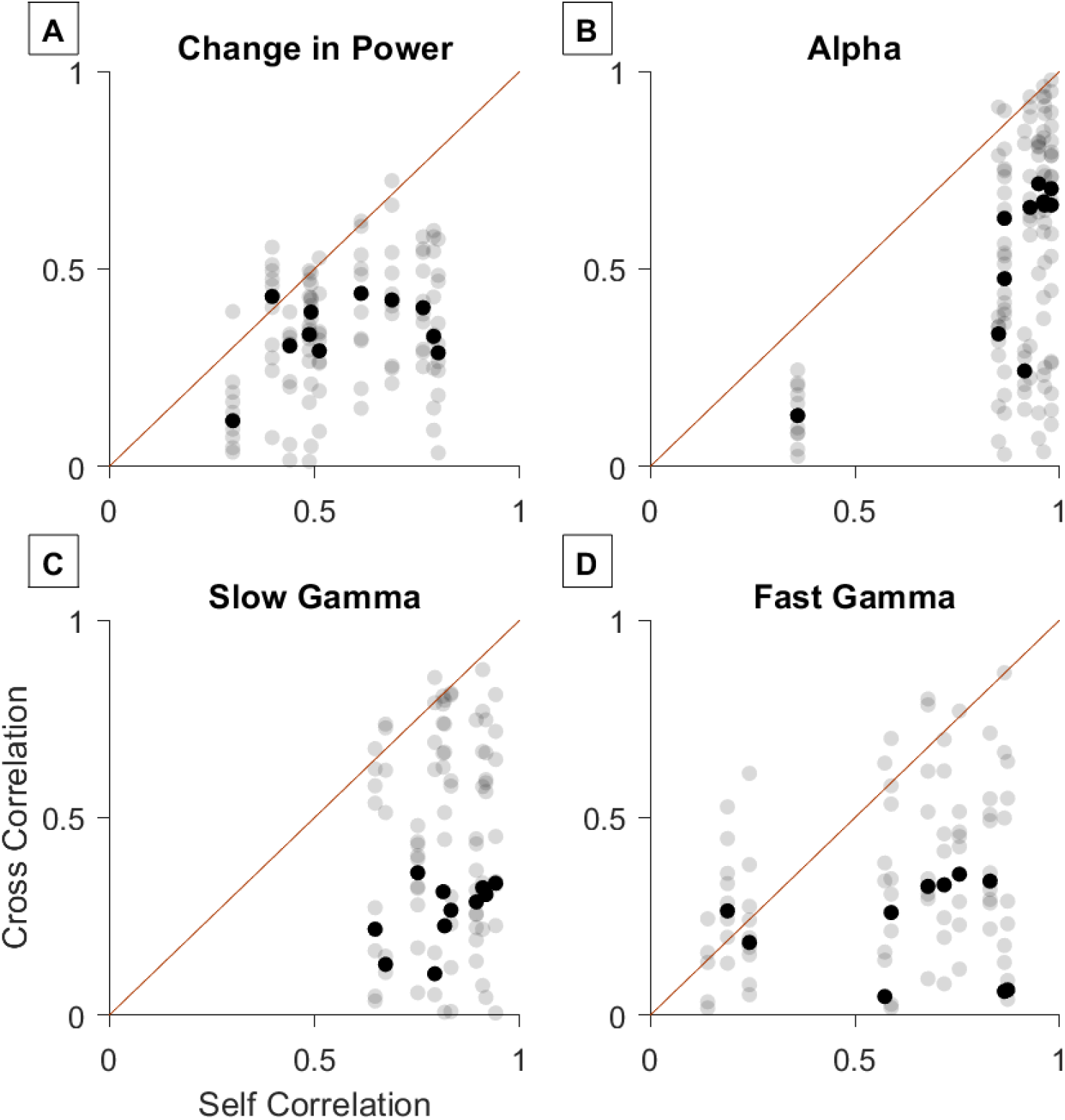
Comparison of self-pair correlations and cross-pair correlations. Self pair correlations (between OpenBCI and Brain Products recordings of same subjects) vs cross pair correlations (between OpenBCI and Brain Products recordings of different subjects) for change in PSD from baseline in stimulus period (**A**) and change in power with time for alpha (**B**), slow gamma (**C**) and fast gamma (**D**) bands. *Red line* indicates x=y line. The *bold black marker* indicates the median of the cross pair correlations for each subject.

## Discussion

Our study is the first one to assess the performance of a low-cost amplifier in the detection of stimulus induced gamma response and compare it with a research grade amplifier while controlling for subjects, electrodes and electrode placement, task performed and subsequent analysis. We showed that a low-cost EEG amplifier like OpenBCI is able to detect gamma response in subjects, and the spectral and temporal profiles from OpenBCI recordings are correlated to that of a research grade amplifier like Brain Products when full screen static gratings are presented as stimuli. However, the change in power in the gamma band was of a lower amplitude in OpenBCI than Brain Products, especially in the fast gamma band. Correlations between the two recordings of the same subjects (‘self-pair correlations’) were significantly higher than correlations between different subjects (‘cross-pair correlations’) indicating that the distinctiveness in spectral and temporal profiles across different subjects could be captured by these amplifiers.

### Comparisons between low-cost and research-grade amplifiers in previous studies

Previous studies have assessed the performance of low-cost EEG amplifiers in various contexts. While some studies have only assessed the performance of a low-cost amplifier in the absence of a research grade control [22, 38], others have compared their performance with a research grade amplifier but have not controlled for factors like the use of same electrodes, same subjects or downstream analysis to reliably attribute all differences only to the amplifiers [39, 40]. de Vos and colleagues [20] compared the performance of an Emotiv based setup with Brain Products GmbH with the same electrodes in the context of a P300 spelling task in 13 subjects. However, they mainly compared the P300 ERP topographies and spelling task performance (*r >* 0.77), with no frequency domain comparison. In a study done by Frey, 2016 [21], OpenBCI signals were compared to the signals from g.tec g.USBamp amplifier (another research grade amplifier) from one subject in a P300 spelling task and a working memory load task. The study used a custom adapter which enabled simultaneous recording with two amplifiers using the same electrodes in the same recording session. The study reported a high correlation between ERPs and PSDs of the two amplifiers (*r >* 0.99). Their simultaneous recording from the two amplifiers in a single recording session as opposed to our sequential recording sessions and their low sample size (*n* = 1) could be an explanation for the lower ERP correlation values (mean *r* = 0.79) and PSD correlation values we found in our study during baseline period (*r* = 0.93). Also, their spectral analyses were restricted to frequencies *<* 40 Hz, which does not include the mains noise range, potentially contributing to a higher correlation value. They also reported a slight jitter (88 ms) in the ERP of OpenBCI compared to g.tec. While we have also found a small jitter in the ERP of OpenBCI (median lag = 8 ms, Fig 4) compared to Brain Products, we did not correct for it since all our analyses were in the frequency domain. Rashid et. al. [24] reported no significant difference in the power of signals between OpenBCI and NuAmps (another research grade amplifier) in the beta band (12–38 Hz, which includes slow gamma frequencies) in 22 participants, but the task performed was a motor task.

### Comparison in performance of OpenBCI for different frequency bands

Previous studies have shown that in the presence of static, full-field visual gratings, two gamma bands are found in the EEG: a slow gamma band (20–34 Hz) and a fast gamma band (35–66 Hz) [12, 13]. In addition, alpha band is also suppressed. In our study, slow gamma found to be better retrieved by OpenBCI than fast gamma in terms of amplitude of change in band power and correlation of its temporal evolution (Fig 3, 6). This could be due to the contamination of fast gamma with the cross-modulation noise bands (Fig 1, dashed lines and Fig 2C, left). Slow gamma and fast gamma bands were both found to be reduced in Alzheimer’s disease and Mild Cognitive Impairment patients in a study done by Murty and colleagues [14]. Further, previous studies have shown that slow gamma band is more reliable with age than fast gamma [13] and shows more inter-subject variability and better test-retest reliability [27]. This raises the possibility of using OpenBCI to detect slow gamma as a biomarker or screening tool in low resource settings.

## Conclusion

OpenBCI is a low-cost EEG amplifier whose open-hardware nature offers customisability and ease of interface with existing equipment, and its lack of bulk offers mobility which allows extensibility of experiments and usage in natural environments outside of dedicated laboratories. Our study suggests that OpenBCI has potential as a low-cost alternative to traditional research grade amplifiers in the detection of stimulus-induced gamma oscillations.

## Declaration of Interests

The authors declare no competing financial interests

## Author Contributions

S.P and S.R. conceived the idea of research, S.P. collected the data; S.P. and S.R. analyzed the data and wrote the paper.

## Funding Disclosure

This work was supported by Tata Trusts Grant, Wellcome Trust/DBT India Alliance (Senior fellowship IA/S/18/2/504003 to SR), and DBT-IISc Partnership Programme.

---

**Figure S1.**
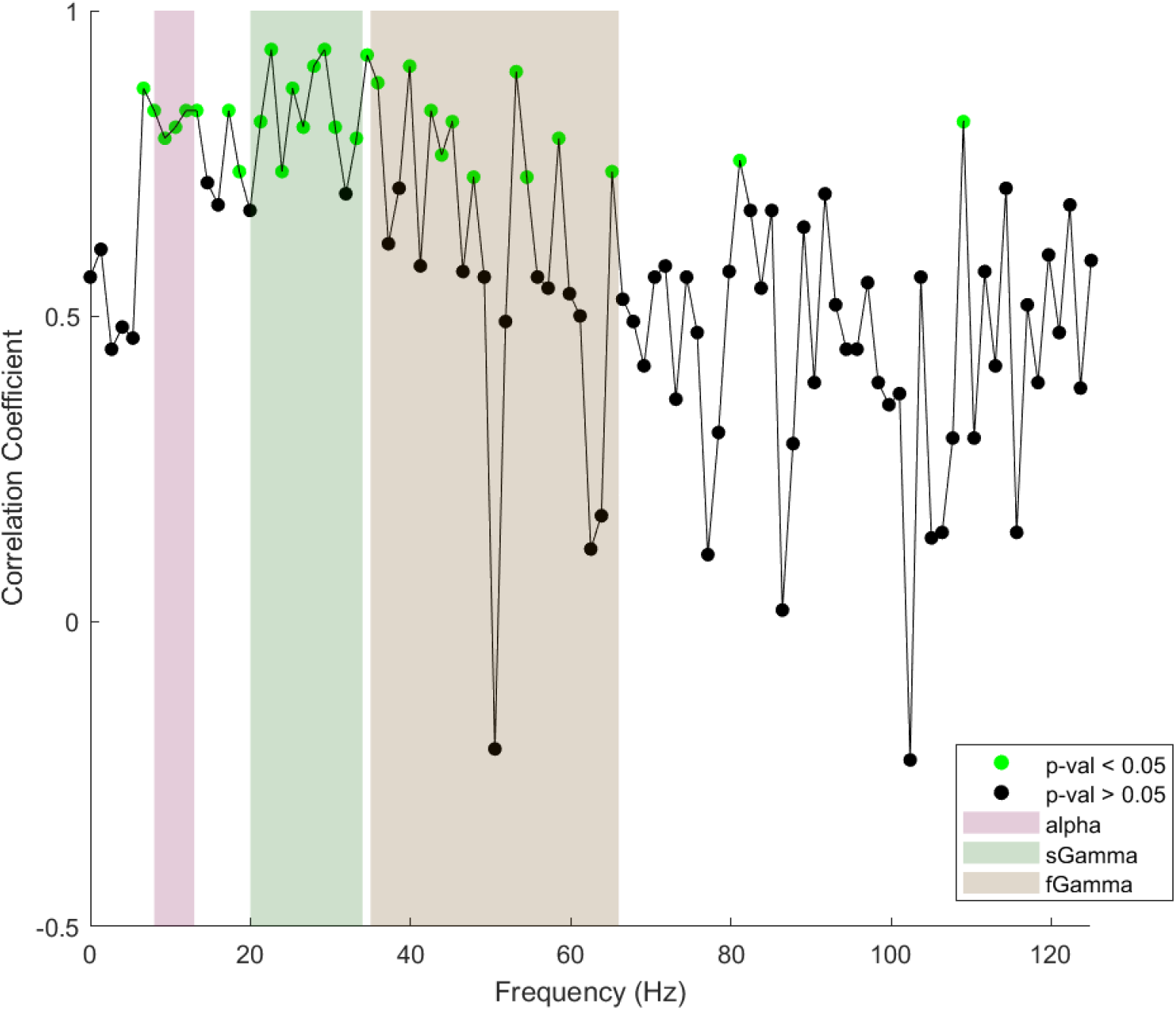
Frequency wise Spearman correlation of power recorded using the two amplifiers across all subjects, plotted across frequency. *Black circles* are for correlation values whose p-values calculated with permutation test are more than 0.05 and *green circles* are for correlation values whose p-values are less than 0.05. False Discovery Rate of p-values is controlled using Benjamini and Hochberg (1995) procedure.

## Notes

https://github.com/FlyingFordAnglia/OpenBCIGammaProject

### Competing Interest Statement

The authors have declared no competing interest.

